# Identification of French Guiana anopheline mosquitoes by MALDI-TOF MS profiling using protein signatures from two body parts

**DOI:** 10.1101/2020.05.22.110452

**Authors:** Sébastien Briolant, Monique de Melo Costa, Christophe Nguyen, Isabelle Dusfour, Vincent Pommier de Santi, Romain Girod, Lionel Almeras

## Abstract

In French Guiana, the malaria, a parasitic infection transmitted by *Anopheline* mosquitoes, remains a disease of public health importance. To prevent malaria transmission, the main effective way remains *Anopheles* control. For an effective control, accurate *Anopheles* species identification is indispensable to distinguish malaria vectors from non-vectors. Although, morphological and molecular methods are largely used, an innovative tool, based on protein pattern comparisons, the Matrix Assisted Laser Desorption / Ionization Time-of-Flight Mass Spectrometry (MALDI-TOF MS) profiling, emerged this last decade for arthropod identification. However, the limited mosquito fauna diversity of reference MS spectra remains one of the main drawback for it large usage. The aim of the present study was then to create and to share reference MS spectra for the identification of French Guiana *Anopheline* species. A total of eight distinct *Anopheles* species, among which four are malaria vectors, were collected in 6 areas. To improve *Anopheles* identification, two body parts, legs and thoraxes, were independently submitted to MS for the creation of respective reference MS spectra database (DB). This study underlined that double checking by MS enhanced the *Anopheles* identification confidence and rate of reliable classification. The sharing of this reference MS spectra DB should made easier *Anopheles* species monitoring in endemic malaria area to help malaria vector control or elimination programs.

## Introduction

Since 2005, malaria cases declined significantly in French Guiana, an oversea territory of France located in South-America. The number of diagnosed cases has decreased from 4,479 cases in 2005 to 597 cases in 2017 [1]. However, the disease is still endemic in the inland forested areas, especially in illegal gold mining areas and so remains of public health importance [2–4]. Majority of the cases are caused by *Plasmodium vivax* (89%, p=531/597), followed by *P. falciparum* (11%, p=66/597).

*Anopheles* mosquitoes are known for their role in transmitting malaria. Historically, thirty-three mosquitoes from the genus *Anopheles* have been reported in French Guiana [5]. Members in the subgenus *Anopheles* and *Nyssorhynchus* have been implicated in malaria transmission in French Guiana. *Anopheles darlingi* is the recognized primary vector in the territory [6–9]. Recently, *An. nuneztovari sl, An. oswaldoi sl, An. intermedius, An. marajoara* and *An. ininii* were found naturally infected with *Plasmodium* sporozoites and were suspected to be secondary vectors [2,3,10]. Malaria transmission is further complicated as some of these secondary vectors belong to species complexes, characterized by similar morphological characteristics, such as *An. oswaldoi* [11], *An. marajoara* [12] and *An. nuneztovari* [13].

A rapid and accurate identification of *Anopheles* species is then critical when designing malaria vector control strategies which should be species-specific to be effective. The most common method for mosquito species identification remains the utilization of morphological criteria [14]. However, morphological identification is skill dependent requiring entomological expertise. The correct species assignation could also be compromised for damaged specimens with the loss of determinant characters. Moreover, the description of morphological characteristic variations between intact adult conspecific specimens underlined that correct mosquito species classification could be held only by experienced mosquito taxonomists [15]. Additionally, the availability of mosquito taxonomic keys is a cornerstone for identification success. Unfortunately, morphological keys are still missing for numerous *Anopheles* species, notably for closely related-species (cryptic and complex species).

The emergence of molecular biology approaches in the 2000’s has solved some long-standing taxonomic questions [16]. However, the choice of the target gene sequence for accurate mosquito identification could be complex. In South America, *Anopheles* species were classified by using different target genes: either 18S rRNA [17], either the second internal transcribed spacer (ITS2) [18] or cytochrome c oxidase I (COI) [19]. The absence of consensus for mosquito species identification complicate studies comparison. The development of the Barcode of Life Data (BOLD) system contributed to standardize gene sequencing for organism identification [20]. Nevertheless, for some cryptic mosquito species (eg, *An. gambiae* complex), the sequencing of COI could be insufficient to classify them unambiguously [21] and the sequencing of a second gene could be required [22]. For instance, some *Anopheles* and *Culex* sibling species could not be distinguished using uniquely the mitochondrial COI barcode [23,24]. Moreover, despite the large advances of this strategy this last decade, notably by shortening experiment duration and costs of reagents, gene sequencing remains time-consuming and expansive [25]. The development of a quick and low cost approach for mosquito monitoring with elevate rate of reliable identification is always in high demand.

The MALDI-TOF MS profiling have recently demonstrated its performance for reliable arthropod identification [25], including mosquitoes at adult [26] and immature stages [27,28]. At adult stage, for specimen identification by MS, different mosquito body parts were selected such as the cephalothorax [29,30] or legs, but this last compartment remains the more frequently used [26,31–33]. The recent successful identification of mosquitoes at immature stages by MALDI-TOF MS profiling validated the efficiency of this MS strategy for field monitoring of *Culicidae* [19]. The regent low costs, the rapidity and technical simplicity of protocols participated to the success of this approach. However, conversely to molecular analyses, MS protein profiles from conspecific specimens could vary according to several factors such as sample storing mode, developmental stage, homogenization mode or body part used [25,34]. To overcome these limitations, standardized protocols were established for some arthropod families [35] including mosquitoes [36]. The standardization of the protocols facilitated result comparisons and reference MS spectra sharing.

The main problem with legs is that they are breakable. The loss of one to all mosquito legs during trapping and/or storing is not infrequent, which could compromise specimen identification by MS. The selection of a second body part, the mosquito thorax, revealed that it could also generate mosquito species-specific MS but distinct from legs of conspecific specimens [37]. This last study underlined that the query of these two body parts against the reference MS spectra database (DB) improved mosquito species identification with accuracy and confidence. This pioneering study assessed to distinct 7 mosquito species from 4 genera living in sympatry in Guadeloupe Island [38].

The aim of the present study was to assess whether the submission of two body part could improve *Anopheles* identification from French Guiana. The creation of a reference MS spectra DB should made easier *Anopheles* species monitoring in endemic malaria area to help malaria vector control or elimination programs.

## Methods

### Mosquito collection and dissection

*Anopheles* adult female mosquitoes were selected from field mosquito collections done in 6 distinct sites from French Guiana, during entomological surveys, using different collection methods, over different sampling periods (Figure 1) [3,39–41]. After collections, mosquitoes were sorted by genera and *Anopheles* mosquitoes were morphologically identified under a binocular loupe at a magnification of ×56 (Leica M80, Leica, Nanterre, France) using standard taxonomic keys for the region (Floch and Abonnenc 1951, Forattini 1962, Faran and Linthicum 1981). *Anopheles* specimens were then stored individually at −20°C. According to their availability, one to 22 specimens per *Anopheles* species were selected for molecular and MS analyses (Table 1). Legs and thoraxes from mosquitoes were dissected for MALDI-TOF MS analysis as previously described [38]. The abdomens, wings and heads were kept for molecular analyses.

**Table 1.**
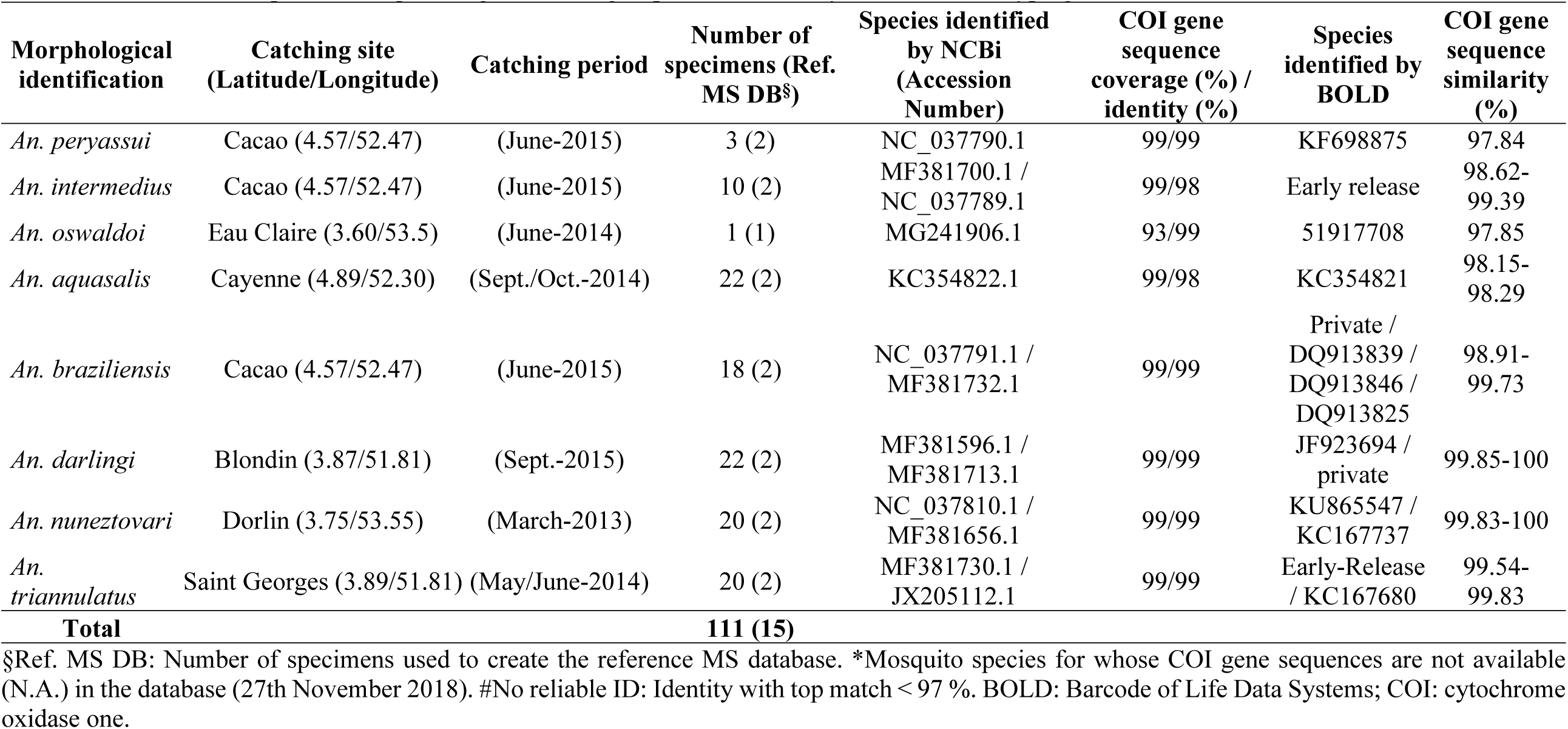
Overview of *Anopheles* mosquito origins and subgroup identification by COI molecular typing

| Morphological identification | Catching site (Latitude/Longitude) | Catching period | Number of specimens (Ref. MS DB§) | Species identified by NCBI (Accession Number) | COI gene sequence coverage (%) / identity (%) | Species identified by BOLD | COI gene sequence similarity (%) |
| --- | --- | --- | --- | --- | --- | --- | --- |
| *An. peryassui* | Cacao (4.57/52.47) | (June-2015) | 3 (2) | NC_037790.1 | 99/99 | KF698875 | 97.84 |
| *An. intermedius* | Cacao (4.57/52.47) | (June-2015) | 10 (2) | MF381700.1 / NC_037789.1 | 99/98 | Early release | 98.62-99.39 |
| *An. oswaldoi* | Eau Claire (3.60/53.5) | (June-2014) | 1 (1) | MG241906.1 | 93/99 | 51917708 | 97.85 |
| *An. aquasalis* | Cayenne (4.89/52.30) | (Sept./Oct.-2014) | 22 (2) | KC354822.1 | 99/98 | KC354821 | 98.15-98.29 |
| *An. braziliensis* | Cacao (4.57/52.47) | (June-2015) | 18 (2) | NC_037791.1 / MF381732.1 | 99/99 | Private / DQ913839 / DQ913846 / DQ913825 | 98.91-99.73 |
| *An. darlingi* | Blondin (3.87/51.81) | (Sept.-2015) | 22 (2) | MF381596.1 / MF381713.1 | 99/99 | JF923694 / private | 99.85-100 |
| *An. nuneztovari* | Dorlin (3.75/53.55) | (March-2013) | 20 (2) | NC_037810.1 / MF381656.1 | 99/99 | KU865547 / KC167737 | 99.83-100 |
| *An. triannulatus* | Saint Georges (3.89/51.81) | (May/June-2014) | 20 (2) | MF381730.1 / JX205112.1 | 99/99 | Early-Release / KC167680 | 99.54-99.83 |
| **Total** |  |  | **111 (15)** |  |  |  |  |

§Ref. MS DB: Number of specimens used to create the reference MS database. \*Mosquito species for whose COI gene sequences are not available (N.A.) in the database (27th November 2018). #No reliable ID: Identity with top match < 97 %. BOLD: Barcode of Life Data Systems; COI: cytochrome oxidase one.

**Figure 1.**
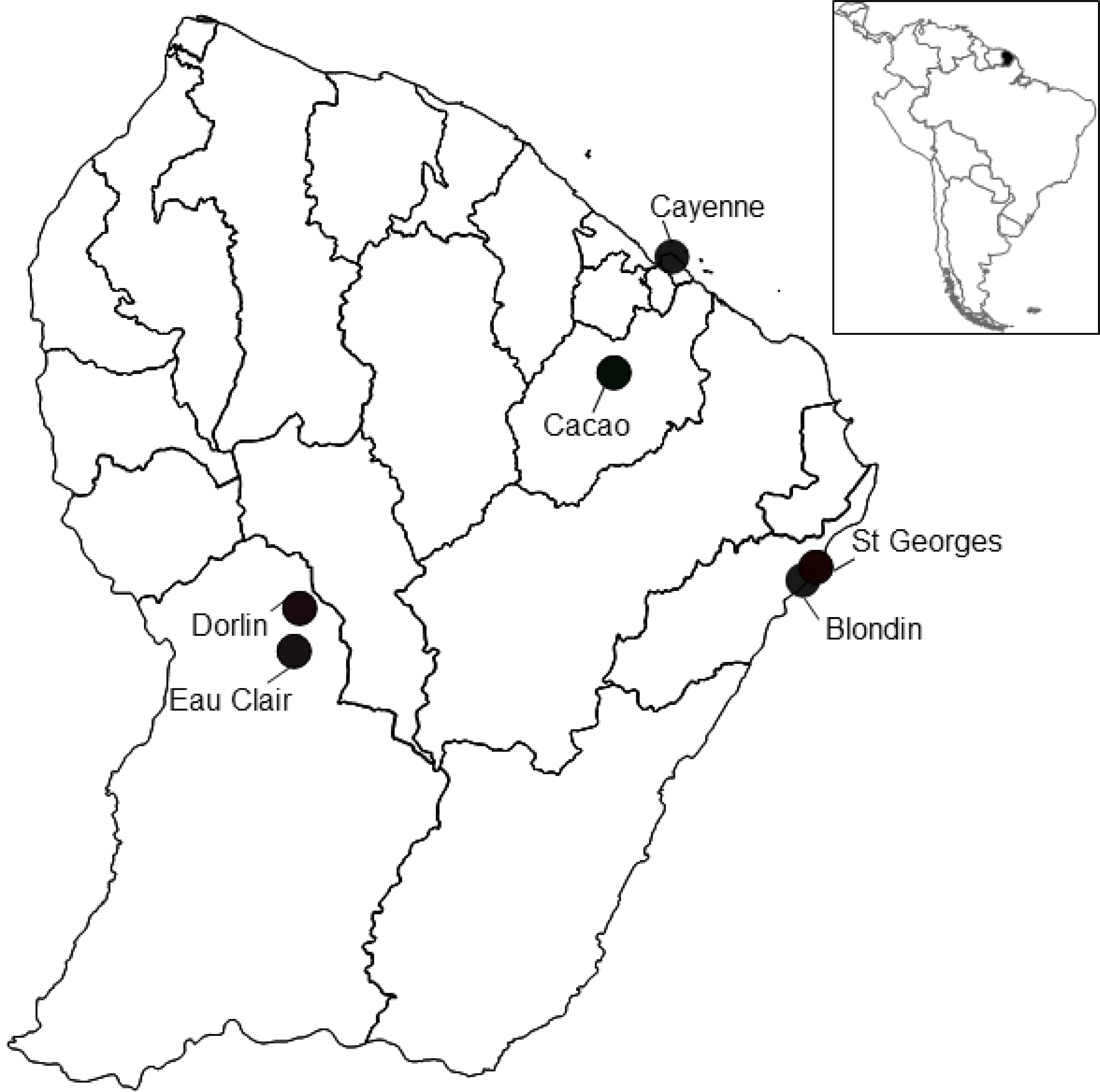
Map of mosquito collection sites in French Guiana. The different sampling sites are indicated by circles.

### Molecular identification of mosquitoes

DNA was individually extracted from the head and abdomen of all mosquito specimens (n=112) using the QIAamp DNA tissue extraction kit (Qiagen, Hilden, Germany) according to the manufacturer’s instructions. Molecular identification of mosquito at the species level was performed by sequencing the PCR product of a fragment of the cytochrome c oxidase I gene (*COI*) (LCO1490 (forward): 5’-GGTCAACAAATCATAAAGATATTGG-3’; HC02198 (reverse): 5’-TAAACTTCAGGGTGACCAAAAAATCA-3’) as previously described [19,42]. The sequences were assembled and analyzed using the Molecular Evolutionary Genetics Analysis (MEGA) software version 7.0 and BioEdit Sequence alignment editor software version 7.2.6.0. All sequences were blasted against GenBank (http://blast.ncbi.nlm.nih.gov/Blast.cgi) and against the Barcode of Life Data Systems (BOLD; http://www.barcodinglife.org; [20]) to assign unknown *COI* sequences to mosquito species.

### Sample homogenization and MALDI-TOF MS analysis

Each dissected compartment (legs and thoraxes) was homogenized individually 3 x 1 minute at 30 Hertz using TissueLyser (Qiagen) and glass beads (#11079110, BioSpec Products, Bartlesville, OK, US) in a homogenization buffer composed of a mix (50/50) of 70% (v/v) formic acid (Sigma) and 50% (v/v) acetonitrile (Fluka, Buchs, Switzerland) according to the standardized automated setting as described previously [43]. After sample homogenization, a quick spin centrifugation at 200 g for 1 min was then performed and 1 µL of the supernatant of each sample was spotted on the MALDI-TOF steel target plate in quadruplicate (Bruker Daltonics, Wissembourg, France). After air-drying, 1 µL of matrix solution composed of saturated α-cyano-4-hydroxycinnamic acid (Sigma, Lyon, France), 50% (v/v) acetonitrile, 2.5% (v/v) trifluoroacetic acid (Aldrich, Dorset, UK) and HPLC-grade water was added. To control matrix quality (i.e. absence of MS peaks due to matrix buffer impurities) and MALDI-TOF apparatus performance, matrix solution was loaded in duplicate onto each MALDI-TOF plate alone and with a bacterial test standard (Bruker Bacterial Test Standard, ref: #8255343).

### MALDI-TOF MS parameters

Protein mass profiles were obtained using a Microflex LT MALDI-TOF Mass Spectrometer (Bruker Daltonics, Germany), with detection in the linear positive-ion mode at a laser frequency of 50 Hz within a mass range of 2-20 kDa. The setting parameters of the MALDI-TOF MS apparatus were identical to those previously used [44].

### MS spectra analysis

MS spectra profiles were firstly controlled visually with flexAnalysis v3.3 software (Bruker Daltonics). MS spectra were then exported to ClinProTools v2.2 and MALDI-Biotyper v3.0. (Bruker Daltonics) for data processing (smoothing, baseline subtraction, peak picking). MS spectra reproducibility was assessed by the comparison of the average spectral profiles (MSP, Main Spectrum Profile) obtained from the four spots for each specimen according to body part with MALDI-Biotyper v3.0 software (Bruker Daltonics). MS spectra reproducibility and specificity taking into account mosquito body part were objectified using cluster analyses. Cluster analyses (MSP dendrogram) were performed based on comparison of the MSP given by MALDI-Biotyper v3.0. software and clustered according to protein mass profile (i.e. their mass signals and intensities). In addition, to visualize MS spectra distribution according to body part, principal component analysis (PCA) from ClinProTools v2.2 software were performed for each species.

The top-10 and top-5 of the most intense MS peaks per mosquito species and per body-part were analyzed with ClinProTools software to estimate their performance to discriminate the *Anopheles* species for each body-part. The parameter settings in ClinProTools software for spectrum preparation were as follows: a resolution of 300; a noise threshold of 2.00; a maximum peak shift of 800 ppm and a match to calibrating agent peaks of 10%. Peak calculation and selection were performed on individual spectra with a signal-to-noise threshold of 2.00 and an aggregation of 800 ppm. Based on the peak list obtained for each body part per species, the top-10 and top-5 of the most intense m/z peaks were selected to include them into the genetic algorithm (GA) model. The selected peaks by the operator gave a recognition capability (RC) value together with the highest cross-validation (CV) value. The presence or absence of all discriminating peak masses generated by the GA model was controlled by comparing the average spectra from each species per body-part.

### Database creation and blind tests

The reference MS spectra were created using spectra from legs and thorax of two specimens per species when available using MALDI-Biotyper software v3.0. (Bruker Daltonics) [26]. MS spectra were created with an unbiased algorithm using information on the peak position, intensity and frequency. MS spectra from mosquito legs and thoraxes were tested against the in-house MS reference spectra DB, including already legs and thoraxes reference MS spectra from eight distinct mosquito species, but none from the *Anopheles* genus [38]. The reliability of species identification was estimated using the log score values (LSVs) obtained from the MALDI Biotyper software v.3.0, which ranged from 0 to 3. According to previous studies [26,44], LSVs greater than 1.8 were considered reliable for species identification. Data were analyzed by using GraphPad Prism software version 5.01 (GraphPad, San Diego, CA, USA).

### Phylogenetic analyses

After gene sequences alignment with the Clustal ω2 algorithm in the MEGA 7.0 software, a maximum likelihood tree based on the COI gene were constructed using the MEGA 7.0 software [45]. The tree with the highest log likelihood was kept. The tree is drawn to scale, with branch lengths measured in the number of substitutions per site. Support for internal nodes was estimated using the nonparametric bootstrap method with 1000 replications.

## Results

### Morphological identification and molecular validation

Among the mosquitoes captured in the 6 distinct sites from French Guiana (Figure 1), uniquely anopheline specimens were selected. These mosquitoes were classified morphologically into eight distinct species, six from the *Nyssorhynchus* subgenus (*An. aquasalis, An. braziliensis, An. darlingi, An. nuneztovari sl, An. triannulatus sl, An. oswaldoi sl*) and two from the *Anopheles* subgenus (*An. intermedius, An. peryassui*) (Table 1). According to their availability, one to 22 specimens per species were included in the present study. A total of 111 *Anopheles* specimens were selected. The *COI* gene sequencing of all specimens was done to valid morphological identification. *COI* gene sequences were queried against GenBank (NCBI) and the Barcode of Life Data (BOLD) Systems. The query of *COI* gene sequences allowed to obtain reliable mosquito species identification for all samples with identity ranges of 98-99% against GenBank and 98.15-100% against BOLD databases (Table 1). Concordant mosquito species identification were obtained between the two molecular DBs. The *COI* gene sequencing corroborated morphological classifications, at the exception of one mosquito. It was morphologically classified as *An. peryassui* but molecular analysis revealed that it was identified as *An. intermedius*. The phylogenetic analysis was done with the *COI* gene sequences of the 15 mosquito specimens selected for MS reference creation (Additional file 1). Mosquitoes belonging to the same subgenus clustered together.

### Reproducible and specific MS spectra from both mosquito body parts

MS profiles of high intensity (>2000 a.u.) were obtained for legs (Figure 2A) and thoraxes (Figure 2B) from each of the 111 mosquitoes submitted to MALDI-TOF MS. Visual reproducible MS spectra were obtained for specimens of the same species according to body part (Figure 2). To evaluate the reproducibility and specificity of MS spectra from legs and thoraxes according to species, cluster analyses were performed. Two specimens per species were used for MSP dendrogram creation, at the exception of *An. oswaldoi* for which only one specimen was available. The clustering of specimens from the same species on the same branch and the absence of species intertwining underlined the reproducibility and specificity of the protein profiles for each *Anopheles* species for legs (Figure 3A) and thoraxes (Figure 3B). Interestingly, *Anopheles* species ordination from MSP dendrograms was not similar between legs and thoraxes of paired species (Figure 3). Nevertheless, *Anopheles* species from the same subgenus clustered on the same branch on both MSP dendrograms. To visualize specificity of MS spectra according body part per *Anopheles* species, PCAs were performed. PCAs revealed a clear separation of the dots corresponding to MS spectra from the legs and thoraxes, confirming a specificity of MS profiles between these two body parts for the seven *Anopheles* species tested (Additional file 2).

**Figure 2.**
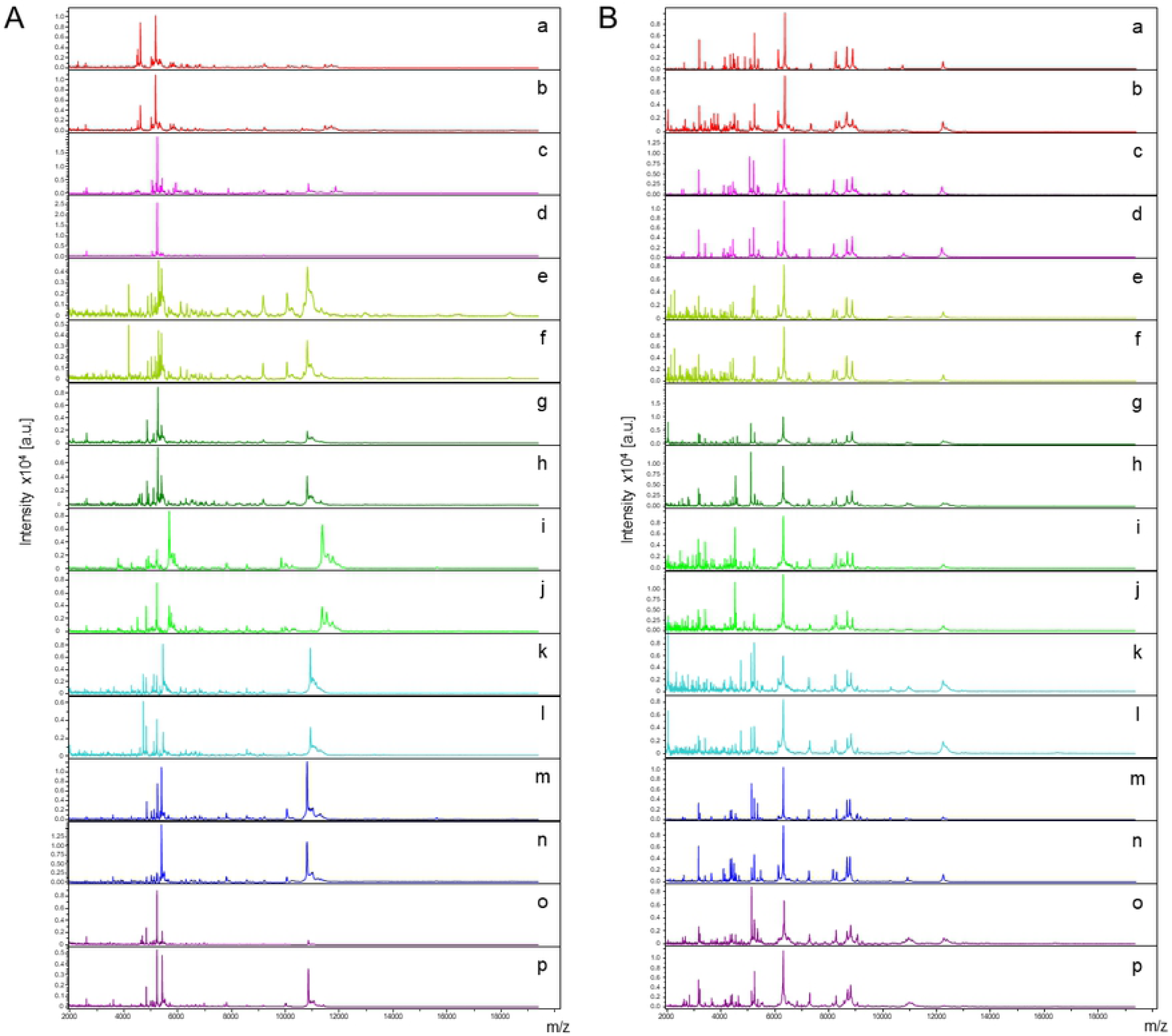
Comparison of MALDI-TOF MS spectra from legs (A) and thoraxes (B) of *Anopheles* mosquitoes. Representative MS spectra of *An. peryassui* (a, b), *An. intermedius* (c, d), *An. oswaldoi* (e, f), *An. aquasalis* (g, h), *An. braziliensis* (i, j), *An. darlingi* (k, l), *An. nuneztovari* (m, n), and *An. triannulatus* (o, p) are shown. MS spectra from two distinct specimens per species were selected, excepted for *An. oswaldoi*. As only one specimen was available for this species, MS spectra from biological replicates were presented. a.u., arbitrary units; m/z, mass-to-charge ratio.

**Figure 3.**
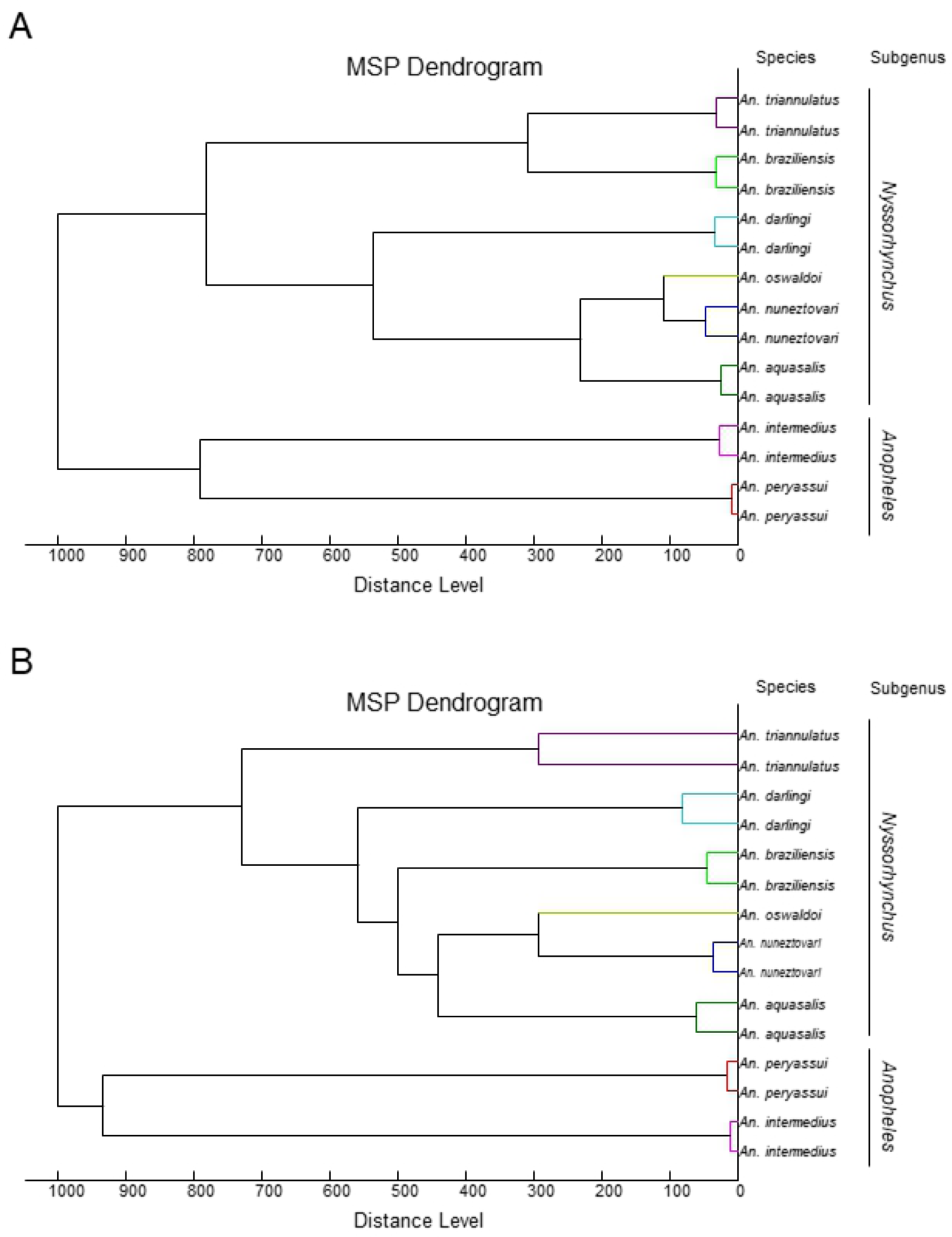
MSP dendrogram of MALDI-TOF MS spectra from legs (A) and thoraxes (B) of *Anopheles* mosquitoes. Two specimens per species were used to construct the dendrogram, at the exception of *An. oswaldoi*, for which only one specimen was available. The dendrogram was created using Biotyper v3.0 software and distance units correspond to the relative similarity of MS spectra. The *Anopheles* and *Nyssorhynchus* subgenus were indicated at the right part.

As correct specimen species classification relies mainly on the intensity of resulting MS spectra, we assessed whether the most intense mass peaks from legs and thoraxes per mosquito species could be enough to distinct these *Anopheles* species, at the exception of *An. oswaldoi* for which only one specimen was available. The selection of the top-ten and top-five mass peak lists per species conducted to a total of 41 and 27 MS peaks for legs and 39 and 24 for thoraxes, respectively (Additional files S3 and S4). These MS peak lists were included in the genetic algorithm (GA) model from ClinProTools 2.2 software. The combination of the presence/absence of these top-ten and top-five mass peak lists per *Anopheles* species displayed, respectively, RC values of 99.6% and 97.0% and CV values of 97.9% and 98.3% for MS spectra from legs. For MS spectra from thoraxes, RC values of 99.4% and 97.5% and CV values of 99.8% and 99.6% were obtained for the top-ten and top-five selected mass peak lists, respectively.

### MS reference spectra database creation and validation step

MS spectra of legs and thoraxes from the 15 specimens used for MSP dendrogram analysis (Table 1), validated morphologically and molecularly, were added to our MS spectra database (DB) including already legs and thoraxes reference MS spectra from eight distinct mosquito species [38]. The legs and thoraxes MS spectra from the 96 remaining specimens were queried against this upgraded DB. Interestingly, 100% of the identification results were concordant between paired MS spectra from legs and thoraxes. Among them, the MS identification of 95 specimens corroborated morphological results. One specimen, classified as *An. peryassui* based on morphological criteria was identified as *An. intermedius* based on MS tool with LSVs of 2.40 and 2.49 for legs and thorax, respectively, confirming the results of *COI* gene sequencing. The LSVs ranged from 1.84 to 2.56 for legs and from 1.60 to 2.61 for thoraxes (Additional file S5). As a threshold LSV upper than 1.8 is require for reliable identification [26,43], correct classification could be considered for 100% (96/96) of MS spectra from legs and 96.9% (3/96) from thoraxes. However, if we considered the LSV results from paired-samples per specimen, 100% of the mosquitoes tested, succeeded to obtain a LSV upper than 1.8 for at least one body-part (Figure 4). Interestingly, an increase of the LSV cut-off at 2.0, which improves identification confidence, revealed that 95.8% (92/96) and 88.5% (85/96) of the specimens reached this threshold, based on their legs and thoraxes MS spectra, respectively. However, the rate of at least one body-part from paired-samples per specimen achieving this threshold (LSVs>2.0) remained at 100%.

**Figure 4.**
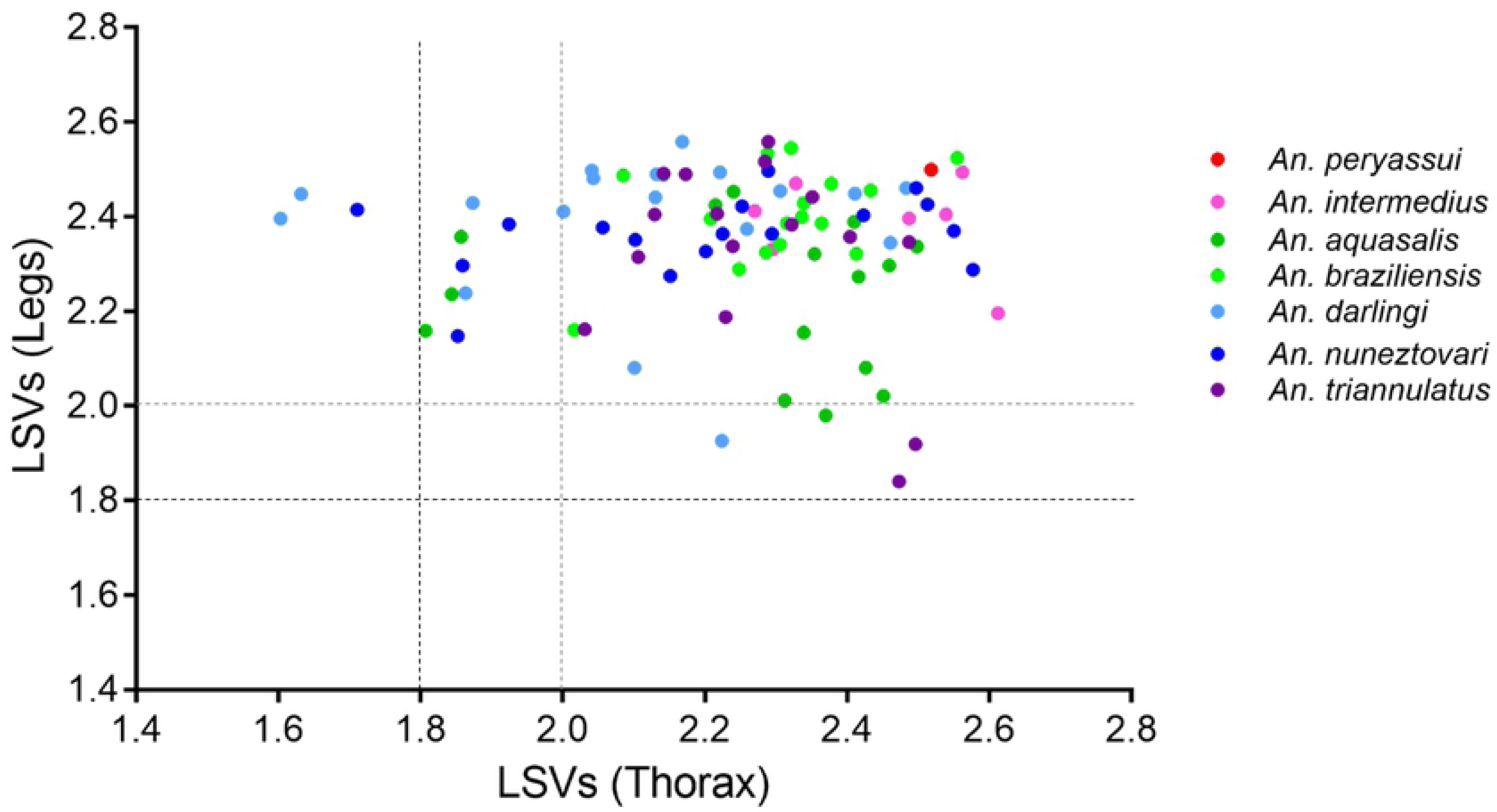
Comparison of paired body parts LSVs from MS spectra of *Anopheles* species. Dashed lines represent the threshold values (black and grey for LSV threshold of 1.8 and 2.0, respectively), for relevant identification. LSV, log score value.

## Discussion

The correct identification of mosquito species is essential for adapted control management, allowing to index circulating species in a given region and, consequently, to estimate vector borne diseases transmission risks. Border regions such as French Guiana may be influenced by their neighboring countries. The legal and illegal flow of human and merchandises through this frontier could induce mosquito vectors migration and conducting to colonization of new areas. Solely, a rapid, accurate and low cost surveillance method of mosquitoes will succeed to improve control measures.

In French Guiana, *An. darlingi* is the main malaria vector transmitting several *Plasmodium* species, such as *P. falciparum* and *P. vivax* [46], this last one representing about 75% of human cases [47]. In addition to *An. darlingi*, other *Anopheles* species were reported to transmit *Plasmodium* pathogens in this area during their blood feeding [10,41]. Moreover these malaria vectors live in sympatry with other anopheline species non-malaria vectors [40]. Indeed, in an effective program aiming to prevent or to eliminate malaria transmission, an accurate identification of *Anopheles* species remains a key factor. The recent repetitive success of the use of MALDI-TOF MS profiling for arthropod identification, including mosquitoes, was applied in the present work to implement our home-made MS reference spectra DB with anopheline specimens from French Guiana and to palliate of the limitations of morphological and molecular analyses [25].

In the present study, we confirmed that legs and thoraxes MS spectra from paired-specimens of the same species were distinct at least for the seven species for which more than one specimen was available. The species-specificity demonstrated for each body-part, underlined that these two compartments could be used independently for mosquito identification. The independent MS submission of several body-part to MALDI-TOF MS for specimen identification, was recently reported for mosquitoes [38] but also for ticks [48,49]. The advantages to test two distinct body parts are the possibility to cross-validate the results and to improve rate of identification confidence and reliability. Effectively, the combination of the results obtained independently by the query against the MS reference DB conducted to concordant results for each body part tested. Here, based on morphological criteria, a specimen was classified as *An. peryassui*. The MS submission of its legs and thorax indicated a matching with *An. intermedius* MS reference spectra for both body parts with high LSVs (>2.40), which was concordant with *COI* gene sequencing. Moreover, if the cut-off threshold of LSV to consider identification as reliable was raised to 2.0, 100% of the specimens succeeded to reach this cut-off with MS spectra from at least one body part. Finally, the concordance of the MS identification results between the two body-parts more the elevate LSVs obtained are complementary data which improve identification confidence. The previous study using two body parts for mosquito identification was done on mosquitoes from distinct genera [38]. Here, the eight mosquito species came all from the *Anopheles* genus including two subgenera, *Nyssorhynchus* and *Anopheles*. The correctness identification of mosquito specimens by MALDI-TOF MS profiling for these close-related *Anopheles* species comforts the accuracy of this innovative tool.

Among the mosquito species included in the MS reference spectra DB, 4 species are malaria vectors (*An. darlingi, An. nuneztovari sl, An. intermedius, An. oswaldoi sl*), for the 4 remaining species their malaria vector competence was not yet demonstrated (*An. aquasalis, An. braziliensis, An. triannulatus sl, An. peryassui*) [5]. Interestingly, malaria vectors and non-vectors are present in each of these two subgenera, underlining the importance to classify them correctly for disease prevention and vector control of species living in sympatric.

The main risks of misidentification or to fail identification by MALDI-TOF MS profiling are generally attributed either to the comprehensiveness of the species included in the MS reference database, or more frequently either to the low intensity of MS spectra. The problem of incomplete reference MS database could be easily solved by performing *COI* gene sequencing of un-matched good quality MS spectra, using the remaining body part of the specimen (ie, head, abdomen or wings). In case of new mosquito species, the addition of respective MS spectra, not yet included in the MS reference DB, could be done. The application of this strategy could resolved step by step MS spectra of good quality failing to find correspondence in the reference MS DB. Concerning the MS spectra un-matched, attributed to the low intensity of protein profiles, this phenomenon is frequently observed for legs MS spectra due to the low protein quantity contained in this compartment [26]. Moreover, as the legs are highly breakable, it is frequent that the loose of one to five legs occurred during specimen collection, transport or storing. This decrease of leg number reduces the success rate of specimen identification [43]. Moreover, for specimens which have lost all legs, their identification become not possible if the identification was based uniquely on this compartment by MS. The creation of reference MS spectra from two distinct mosquito body parts allows to succeed specimen identification by this rapid proteomic approach.

Moreover, in the present work, it was highlighted that the classification of *Anopheles* species could be correctly done using the most intense MS peaks. Effectively, the selection of the top-10 or also the top-5 of the MS peaks possessing the higher intensity appeared sufficiently discriminants to classify correctly these mosquito species. This correct classification is valid for both body-parts. These results underline that a correct identification remain possible with MS spectra of low intensity for which uniquely the most intense MS peaks could be detected.

The comparison of MSP dendrograms between legs and thoraxes revealed that, despite species from the same subgenus were clustered in the same branch, the ordination of the species inside these branches was not similar. This unreproducible classification objectifies that the MS profiles proximity were different between the *Anopheles* species for legs and thorax. These distinct ordination of species based on their MS profiles, should reduce the risk of misidentification by the submission of both body parts. Interestingly, none of these MSP dendrograms proposed a classification comparable with those obtained using *COI* gene sequences for phylogenetic tree construction. The MSP dendrograms are not adapted for phylogenetic analyses as previously reported [50,51].

## Conclusion

Mosquito monitoring with fast, highly reproducible and reliable tools such as MALDI-TOF appears essential in today’s globalization scenario. The specificity of MS protein profiles for mosquito legs and thoraxes confirmed that these two body parts are relevant for specimen identification. Moreover, as the most intense MS peaks were demonstrated to be sufficient for correct classification, the sample which will generate MS spectra of low quality, could anyway identified. The sharing of reference MS spectra is primordial to accelerate the dissemination of this innovative tool for a routine use in mosquito identification contributing to adapt control of vectors.

## List of abbreviations

*sl*: *sensu lato*;
*ss*: *sensu stricto*;
MALDI-TOF MS: Matrix Assisted Laser Desorption/Ionization Time-of-Flight Mass Spectrometry;
PCR: Polymerase Chain Reaction;
CCI: Composite Correlation Index;
LSV: Log Score Value

## Declarations

### Ethics approval and consent to participate

Not applicable.

### Consent for publication

Not applicable.

### Availability of data and materials

The datasets of MS reference spectra added to the MS DB in the current study are freely available and downloadable from the additional file 6.

### Competing interests

The authors declare that they have no competing interests.

### Funding

This work has been supported by the Délégation Générale pour l’Armement (DGA, MoSIS project, Grant no PDH-2-NRBC-2-B-2113).

### Authors’ contributions

Conceived and designed the experiments: LA, SB. Performed the experiments: LA, SB, CNG. Analyzed the data: LA, SB. Contributed reagents/materials/analysis tools: MMC, CNG, VPS, ID, RG. Field collections: SB, VPS, ID, RG. Drafted the paper: LA, SB, MMC. Revised critically the paper: all the authors.

## Acknowledgments

We would like to acknowledge Samuel Vezenegho and Antoine Adde, from Unite d’Entomologie Médicale, Institut Pasteur de la Guyane, Cayenne, French Guiana, for their involvement in sample management. We also acknowledge Albin Fontaine (UPE, IRBA, Marseille) for his help in map building.

## Additional files

**Additional file 1. Unrooted Maximum-Likelihood trees based on the sequences of the COI gene of the 15 specimens included in the MS database.**

**Additional file 2. Principal Component Analysis (PCA) from MS spectra of legs and thoraxes from *Anopheles* mosquitoes.** PCA dimensional image from MS spectra of legs (red dots) and thoraxes (green dots) from *An. intermedius* (A), *An. aquasalis* (B), *An. braziliensis* (C), *An. darlingi* (D), *An. nuneztovari* (E), *An. triannulatus* (F) and *An. peryassui* (G). Respectively, 10, 22, 18, 22, 20, 20 and 3 specimens per species were included. Quadruplicate of each sample per body part were presented.

**Additional file 3. Top-five and -ten mass peak lists per mosquito species using legs as biologic material.**

**Additional file 4. Top-five and -ten mass peak lists per mosquito species using thoraxes as biologic material.**

**Additional file 5. LSVs obtained following homemade MS reference database query with MS spectra of legs (A) and thoraxes (B) from *Anopheles* mosquitoes.** Horizontal dashed lines represent the threshold value for reliable identification (black and grey for LSV threshold of 1.8 and 2.0, respectively). LSVs, log score values; a.u., arbitrary units.

**Additional file 6. Raw MS spectra from legs and thoraxes of mosquitoes added to the MS reference database.** MS spectra were obtained using Microflex LT MALDI-TOF Mass Spectrometer (Bruker Daltonics).

